# MicroRNA-202 safeguards meiotic progression by preventing premature degradation of REC8 mediated by separase

**DOI:** 10.1101/2021.04.14.439735

**Authors:** Jian Chen, Chenxu Gao, Chunwei Zheng, Xiwen Lin, Yan Ning, Longfei Ma, Wei He, Dan Xie, Kui Liu, Chunsheng Han

## Abstract

MicroRNAs (miRNAs) are believed to play important roles in mammalian spermatogenesis but the *in vivo* functions of single miRNAs in this highly complex developmental process remain unclear. Here, we reported that *miR-202*, a member of the *let-7* family, played an important role in mouse spermatogenesis by phenotypic evaluation of *miR-202* knockout (KO) mice. In *miR-202* KO mice, germ cells underwent apoptosis. Multiple processes in meiosis I including synapsis and crossover formation were disrupted, and inter-sister chromatid synapses were detected. More importantly, we found that upon *miR-202* KO, meiotic-specific cohesin protein REC8 was prematurely cleaved by precociously activated separase, whose mRNA was a direct target of *miR-202-3p*. Our findings identify *miR-202* as a novel regulator of meiosis and contribute to the list of miRNAs that play specific and important roles in developmental processes.

## Introduction

Spermatogenesis is a unique cellular developmental process, which involves many cell-type specific events and a large number of cell-type specific genes (1). During meiosis, homologous chromosomes pair and recombine, culminating in the formation of synaptonemal complex and crossovers, and then segregate into daughter cells (2, 3). Meiosis involves the highly complex yet ordered DNA and protein interactions. For example, synaptonemal complex is a huge tripartite protein structure consisting of two lateral elements, of which SYCP2/3 and cohesin are key components, and a central element, which contains proteins such as SYCP1 and SYCE1/2 (4). The timely production and degradation of proteins in such a complex is one important aspect of the regulatory mechanism of meiosis.

Cohesin is a multi-protein complex that plays a role in the cohesion of sister chromatid and the establishment of higher order chromosome architecture in both somatic and germ cells (5). Somatic cohesin has four core subunits: SMC1a, SMC3, RAD21, and SA1 or SA2. RAD21 is a member of the kleisin family proteins that can be cleaved by the protease separase (6). After RAD21 is cleaved by separase at the anaphase of the cell cycle, sister chromatids are segregated into daughter cells (5). Meiotic cohesin contains meiosis-specific subunits such as SMC1β, REC8, RAD21L1 and SA3/STAG3. Both REC8 and RAD21L belong to the kleisin family (5). During meiosis, cohesin serves as the core for the assembly of chromosome axes, which develop into the lateral elements of the synaptonemal complex (5). Cohesin is not only essential for synapsis, but also for homologous chromosomes to align at the equatorial plate at metaphase I when synaptonemal complex is dismantled and crossovers are formed (5, 7-12). RAD21L is removed from chromosomes at the later stages of meiotic prophase I in a protease independent way, while REC8 beyond centromeres is cleaved by separase and removed from the chromosomes at anaphase I (13, 14). The essentiality of cohesin in gametogenesis has been revealed by the infertility and multiple meiotic defects of the gene KO mice of its components (5, 7-12). For example, in the absence of REC8, DNA double strand breaks (DSBs) cannot be repaired, and synapsis occurs between sister chromatids instead of homologous chromosomes (7, 8).

MicroRNAs are small RNAs (∼22 nt), which are fundamental to the developmental, physiological, and disease processes of metazoans as they can direct the posttranscriptional repression of mRNA targets in diverse lineages (15). Global miRNA loss resulted from the KO/mutations of key regulators in miRNA biogenesis induces dramatic phenotypic changes in almost all examined tissues, whereas individual miRNA KO often lacks dramatic phenotypic consequences, implying that miRNAs act in a redundant manner (16). miRNAs are believed to be essential for mammalian spermatogenesis as demonstrated by the infertile phenotypes of the KO mice of genes of key regulators of miRNA biogenesis, such as *Dicer, Drosha*, and *Dgc8* (17-25). However, physiological functions of single miRNA genes in spermatogenesis have not been reported, which may mainly because most miRNA genes are clustered within small regions of the genome.

The intergenic miRNA gene *miR-202* belongs to the highly conserved *let-7* family (26) and generates *miR-202-3p* and *miR-202-5p*, two complementary miRNAs that are highly enriched in mouse testes (27). In the present study, we constructed *miR-202* KO mice by using the CRIPSR-Cas9 technique and found that spermatogenesis in these mice, particularly the prophase of meiosis I, was overtly impaired. Further, we showed that the separase-REC8 pathway was pre-activated in the absence of *miR-202* and separase mRNA was a direct target of *miR-202-3p*. For the first time, we find that a single miRNA gene plays an important role in spermatogenesis and demonstrate the regulation of the meiotic cohesin by a single miRNA. These results add to our understanding of the regulatory mechanism of meiosis.

## Results

### *MiR-202* KO impairs spermatogenesis in mice

We investigated the *in vivo* function of *miR-202* via phenotypic evaluation of *miR-202* KO mice that were constructed with the CRISPR-Cas9 technique. A 155-bp fragment containing the whole transcribed region of *miR-202* was deleted in a male founder mouse (Fig. 1A), and this mutant allele was passed to F3 mice, which were then used for phenotypic evaluation (Fig. S1A-S1C). Neither the male nor the female homozygous KO mice (hereafter KO representing homozygotes) exhibited any overt physical abnormalities compared with their wild-type (WT) littermates (Fig. 1B and S1D). However, the testis weight of the male KO mice, normalized by their body weight, was significantly reduced (Fig. 1B and 1C). The fertility of female KO mice was comparable to that of the female WT mice (Fig. S1E), whereas, the mating of KO males with WT females resulted in significantly lower pregnancy rate and litter size compared with the WT males (Fig. 1D and 1E).

**Fig. 1.**
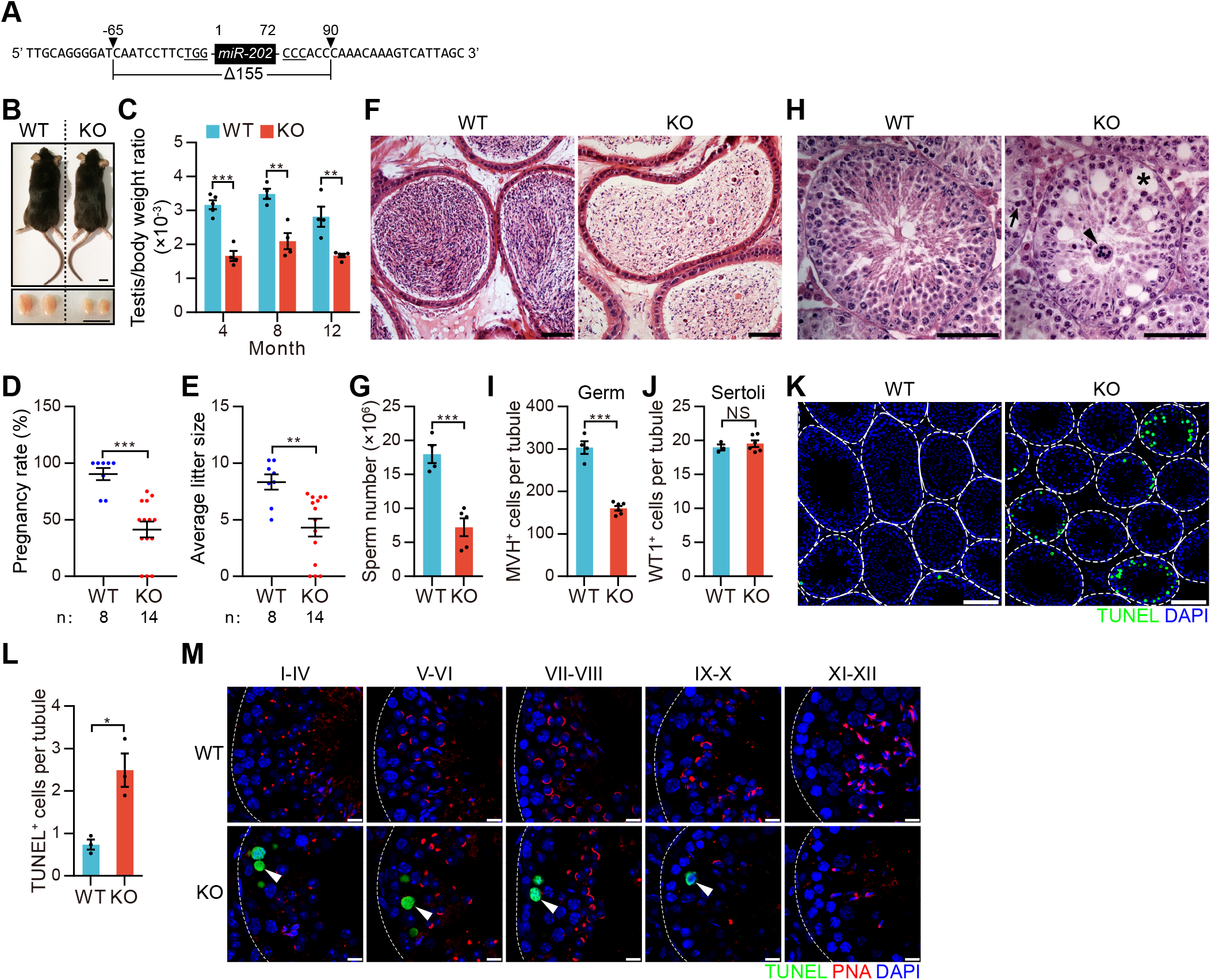
*miR-202* KO impairs spermatogenesis in mice. (**A**) A schematic diagram of the deletion of *miR-202* loci. PAM sites of sgRNAs are underlined. (**B**) Representative images of mice and testes at 4 months of age. (**C**) Quantitative comparison of testis/body weight ratio between WT and KO mice. (**D** and **E**) Pregnancy rate (D) and average litter size (E) of adult WT or KO males. Each male mated with at least three WT females and the plugged females were counted. Each point represents one male. (**F**) HE staining of caudal epididymis sections from WT and KO mice at four months of age. (**G**) Caudal epididymal sperm counts of adult WT and KO mice. (**H**) HE staining of testis sections from WT and KO mice at four months of age. Arrowhead and arrow indicate multinucleated syncytia and an apoptotic cell, respectively. Asterisk marks a vacuole. (**I** and **J**) Quantitative comparison of MVH^+^ (I) or WT1^+^ (J) cells per tubule between adult WT and KO mice. At least 20 tubules for each mouse were counted. (**K** and **L**) TUNEL staining of testis sections from WT and KO mice at four months of age (K) with apoptotic cell number per tubule (L). White dotted line indicates the outline of testis tubules. (**M**) Validation of apoptotic cell type by combining TUNEL assay with PNA staining. Arrowheads indicate apoptotic pachynema based on their more lumenal position within the tubules. White dotted line indicates the outline of testis tubules. All values are shown as mean ± SEM. *p < 0.05, **p < 0.01, ***p < 0.001. NS, not significant. Scale bars: 1cm (B), 100 μm (K), 50 μm (F, H) and 10 μm (M).

The epididymides of the adult male KO mice contained lower number of sperms; however, the morphology of these sperms was normal under the microscope (Fig. 1F, 1G, and S1F). Many vacuoles were observed on the testis sections of the KO mice, and the detached multinucleated syncytia were also observed (Fig. 1H). The number of MVH^+^ germ cells of the KO mice was significantly reduced than that of WT mice, while the number of Sertoli cells marked by the specific expression of the transcription factor Wilm’s Tumor 1 (WT1) was not significantly different from that of the WT mice (Fig. 1I, 1J, S1G, and S1H). The seminiferous tubules of the KO mice contained many apoptotic cells, in which the nuclei were condensed and strongly stained on hematoxylin-eosin (HE) stained sections (Fig. 1H). The apoptosis in KO mice was further confirmed by TUNEL assays (Fig. 1K and 1L). Furthermore, the apoptotic cells were identified to be spermatocytes based on the triple staining of nuclei by 4, 6-diamidino-2-phenylindole (DAPI), the acrosome by peanut agglutinin (PNA), and the apoptotic cells by TUNEL (Fig. 1M) (28). These results indicate that *miR-202* plays an important role in spermatogenesis by sustaining the optimal population and viability of spermatogenic cells.

### Changes in cellular composition and gene expression upon *miR-202* KO revealed by single-cell RNA-seq (scRNA-seq)

As there are all types of testicular cells in KO mice, we next used scRNA-seq analyses to investigate the changes in cellular composition and gene expression upon *miR-202* KO. One testis from each of three adult WT mice and three adult KO mice was collected, and the KO and WT testes were pooled respectively for subsequent analysis (Fig. S2A). A total of 15,310 WT and 12,935 KO cells passed standard quality control and were used for subsequent analysis (Fig. S2B and Table S1). On average, we identified 11,036 and 12,272 UMIs (unique molecular indices), corresponding to 2,896 and 3,074 genes for WT and KO mice, respectively (Fig. S2B and Table S1). Unsupervised clustering of the total 28,245 testicular cells were shown by the Manifold Approximation and Projection (UMAP) analysis plots (29), and 16 major cell types for both WT and KO mice were identified (Fig. 2A and S3A). Cluster identity was assigned based on expression patterns of known marker genes in mouse testes, including spermatogonia (SG; marker: *Uchl1*), spermatocytes (SCs; markers: *Sycp1, Ccna1* and *Spag6*), round spermatids (rSTs; marker: *Fam71b*), elongating spermatids (eSTs; marker: *Prm2*) and Sertoli (marker: *Cst9*) (Fig. 2B and S3A) (30-32). The validity of the clustering was confirmed by the similar UMAP results of our WT cells and the ones of adult C57BL/6J mice published by Ernst *et al*. (Fig. S2C and S2D) (32). As the number of Sertoli cells was not significantly affected by *miR-202* KO (Fig. 1J), we compared the cellular composition of KO and WT testes by using the number of each cell types that was normalized by the number of Sertoli cells. Consistent with the results of testis weight and MVH^+^ cell count, the total number of spermatogenic cells, and the number of meiotic and post-meiotic cells were significantly reduced in KO mice compared with their WT littermates (Fig. 2C and Table S1).

**Fig. 2.**
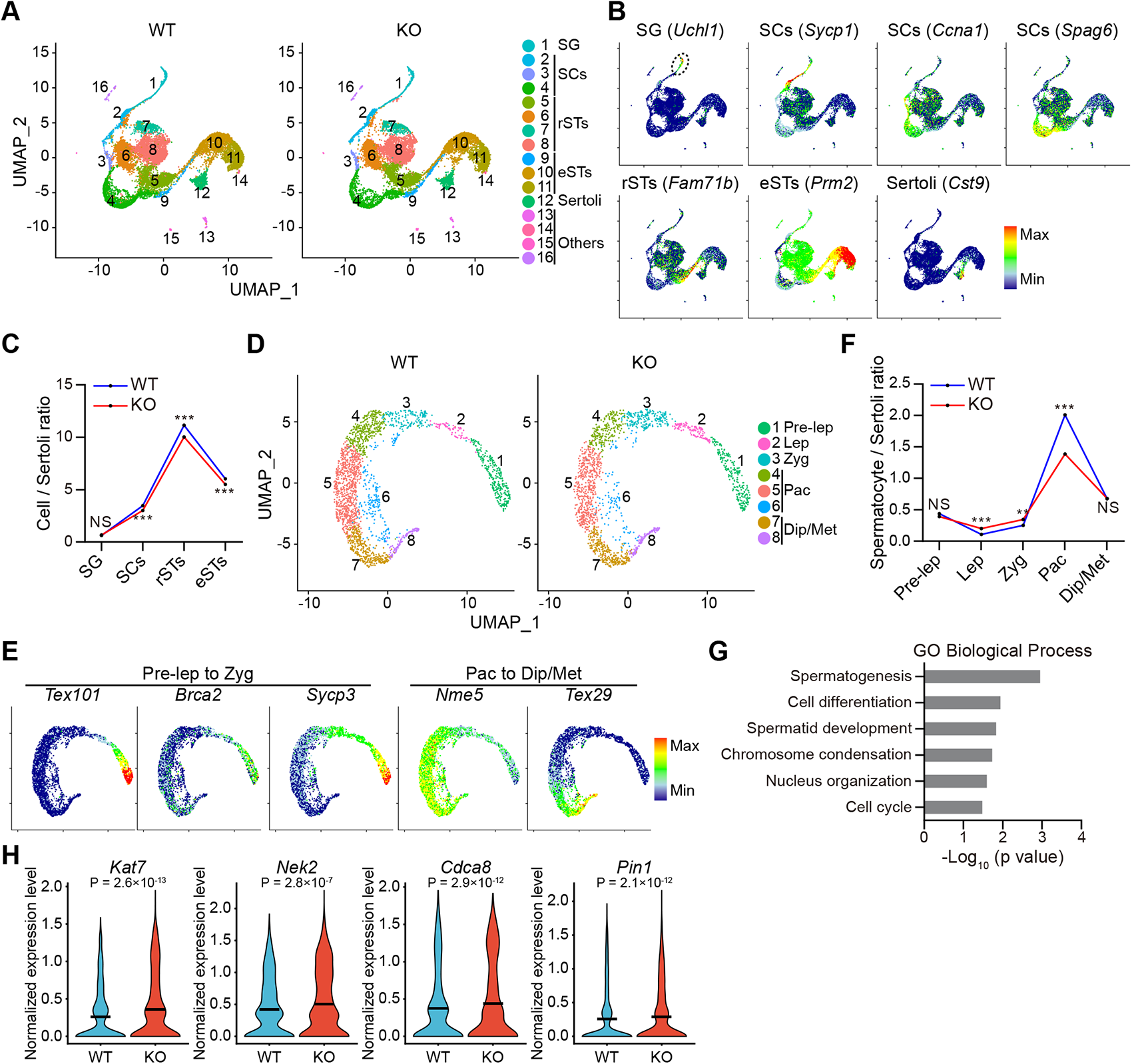
scRNA-seq analysis reveals changes in cellular composition and gene expression upon *miR-202* KO. (**A**) UMAP and clustering analysis of single-cell transcriptome data from adult WT and KO testicular cells. Each dot represents a single-cell and cell clusters are distinguished by colors. (**B**) Gene expression patterns of selected cell marker genes projected on the combined UMAP plot of total cells. (**C**) Cell/Sertoli ratio for each cell type in (A). (**D**) UMAP and re-clustering analysis single-cell transcriptome data from WT and KO SCs of cluster 2, 3 and 4 in (A). Each dot represents a single-cell and cell clusters are distinguished by colors. (**E**) Gene expression patterns of selected cell marker genes projected on the combined UMAP plot of spermatocytes. (**F**) Cell/Sertoli ratio for each cell type in (D). (**G**) Representative GO terms of differentially expressed genes (DEGs) between WT and KO SCs. (**H**) Violin plots of representative DEGs upregulated in KO SCs. The lines represent the mean values. **p < 0.01, ***p < 0.001. NS, not significant.

The 2,351 WT and 1,884 KO SCs were selected for re-clustering analyses (Table S1). Based on the expression patterns of known marker genes for SCs at different stages, the following five types of SCs were identified: pre-leptonema (Pre-lep), leptonema (Lep), zygonema (Zyg), pachynema (Pac), and diplonema/metaphase (Dip/Met) (Fig. 2D, 2E and S3B). Marker genes included *Tex101* (Pre-lep), *Brca2* (Lep), *Sycp3* (Zyg) and *Nme5* (Pac) (32), as well as a marker gene (*Tex29*) expressed in Dip/Met defined by Wang *et al*. (Fig. 2E) (33). Based on the cell numbers normalized by Sertoli cell population, we found that Lep and Zyg increased by 88% and 36%, respectively, while that of Pac decreased by 31% in KO testes compared with the WT ones (Fig. 2F), which represented a blockade from Zyg to Pac.

Next, we performed differentially expressed gene (DEG) analysis on total SCs of WT and KO mice and identified 28 DEGs (P_adj_ < 0.05; Table S2). Gene ontology (GO) analyses showed that these DEGs were enriched in functional annotation terms related to spermatogenesis (spermatogenesis, cell differentiation, spermatid development, chromosome condensation, nucleus organization and cell cycle) (Fig. 2G and Table S2). Several genes that may play a role in spermatogenesis were in the up-regulated gene list (Fig. 2H and Table S2). As a member of the acetyltransferase family, KAT7 (histone acetyltransferase) is required for accurate chromosome segregation by protecting centromeres from the surrounding heterochromatin in mitosis (34). NEK2 (NIMA-related kinase 2) plays an active role in chromatin condensation in mouse pachytene spermatocytes (35) and is critical for proper assembly of the meiotic spindles in mouse oocytes (36). CDCA8 (cell division cycle associated 8) is associated with meiotic spindle assembly and chromosome segregation in human oocytes (37). PIN1 (peptidyl-prolyl cis/trans isomerase) interacts with separase and determines the half-life of enzymatic activity of separase to trigger proper sister chromatid separation in mitosis (38). Together, scRNA-seq results reveals that loss of *miR-202* induces considerable changes in cell composition and transcriptome signatures in SCs.

### Loss of *miR-202* disrupts meiosis I prophase

We next examined whether meiosis is impaired in *miR-202* KO mice. The meiotic prophase of SCs is divided into the Lep, Zyg, Pac, and Dip, which can be distinguished by the co-immunostaining patterns of SYCP3, and the phosphorylated form of histone H2AX (γH2AX), which marks the DSBs generated in Lep/Zyg and the partially synapsed XY chromosomes (also known as XY body or sex body) in Pac/Dip (39). In Lep/Zyg, γH2AX staining was diffuse as a result of the hundreds of DSBs formed in the nucleus, whereas, in Pac/Dip, it was a bright dot representing the sex body (Fig. 3A). We predict that no more than 30% of the tubules of the WT mice might contain both diffuse and dot-like γH2AX signals (2-staining, stages VII-XI) based on the frequencies and the cellular compositions of seminiferous tubules at different stages (40). Indeed, we found that in WT mice, the percentage of the tubules with 2-staining was 26% while that of the tubules with only the dot-like signal (dot-only) was 63%. However, the percentages of these two types of tubules were 42% and 49%, respectively, in KO mice (Fig. 3B and S4). Moreover, the KO testes contained more Lep/Zyg and fewer Pac/Dip as indicated by the γH2AX staining compared with the WT ones (Fig. 3C and S4). These results indicate that *miR-202* KO results in a blockade of meiosis in transition from Zyg to Pac, consistent with our scRNA-seq result (Fig. 2F).

**Fig. 3.**
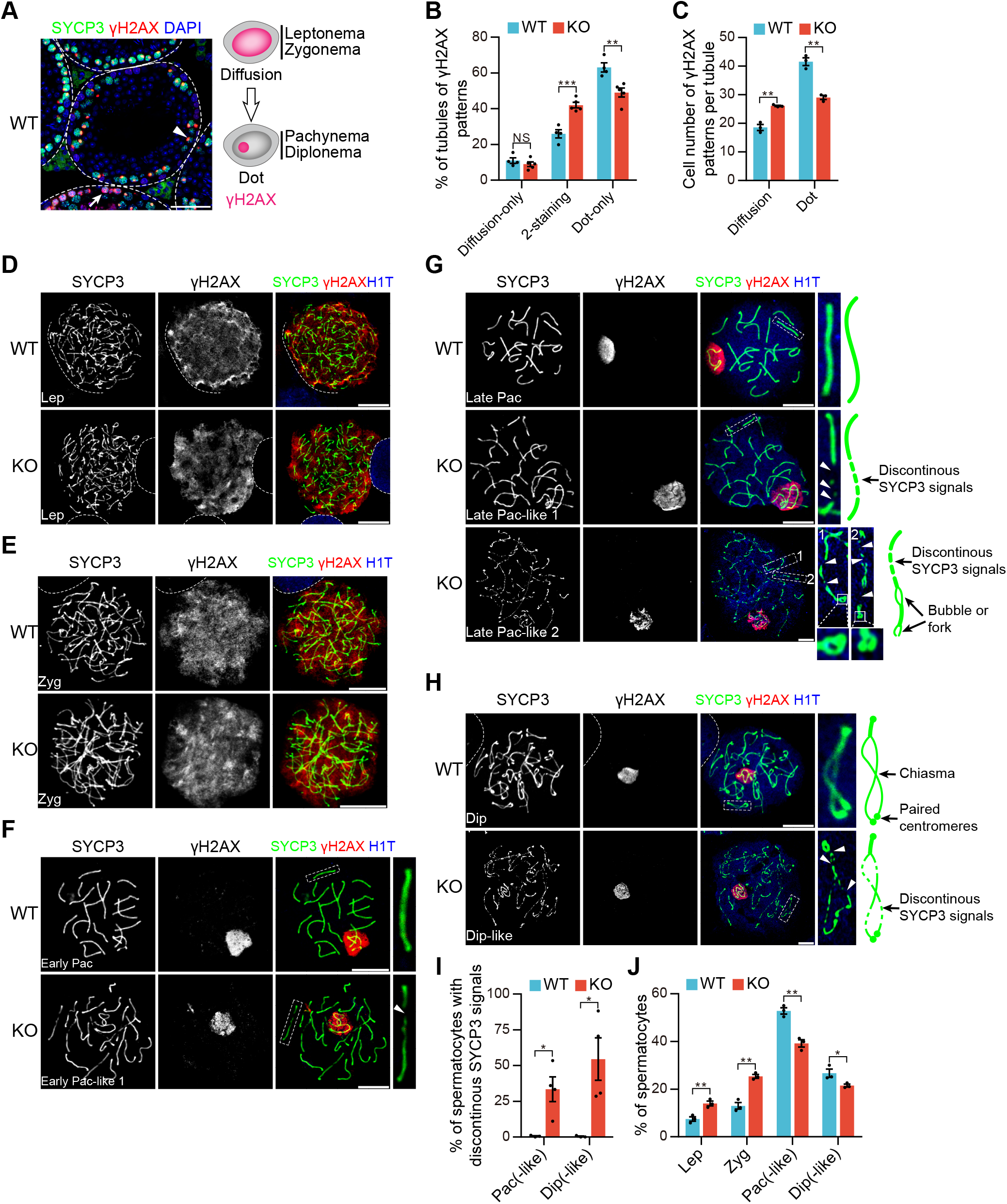
*miR-202* knockout disrupts meiosis I prophase. (**A**) Immunostaining of SYCP3 and γH2AX in sections from adult WT mice. Arrow and arrowhead indicate diffuse and dot-like γH2AX staining, respectively. White dotted line indicates the outline of testis tubules. (**B**) Proportion of tubules of γH2AX expression patterns. At least 100 tubules of each mouse were counted. (**C**) Average number of spermatocytes with γH2AX expression patterns per tubule. At least 50 tubules of each mouse were counted. (**D-H**) Representative nuclear spreads of WT and KO SCs at Lep (D), Zyg (E), early Pac (F), late Pac (G) and Dip (H). Spermatocytes were immunostained for SYCP3, γH2AX, and H1t. Magnified views are indicated by dashed areas. Schematic representation shows homolog characteristic revealed by SYCP3 signals at different backgrounds and stages. Arrowheads indicate bivalent stretches lacking SYCP3 signals in KO. (**I**) Proportion statistics of spermatocytes with discontinuous SYCP3 signals. At least 200 cells of each mouse were counted. (**J**) Meiotic stage frequencies in (D-H). At least 200 cells of each mouse were counted. All values are shown as mean ± SEM. *p < 0.05, **p < 0.01, ***p < 0.001. NS, not significant. Scale bars: 50 μm (A) and 10 μm (D-H).

SCs at different stages of prophase I can be better distinguished by the SYCP3-γH2AX co-immunostaining of surface-spread chromosomes. The testis-specific histone variant H1t is only expressed in mid/late Pac and Dip among all SC (41). Triple staining of SYCP3, γH2AX, and H1t revealed that all types of SCs including late Pac and Dip were present in KO mice (Fig. 3D-3H). No obvious difference in SYCP3 staining between WT and KO Lep/Zyg was detected (Fig. 3D and 3E). However, in KO mice, many Pac/Dip contained chromosome(s) with discontinuous SYCP3 staining indicative of partially formed synaptonemal complex axes and regional asynapsis while such cells were absent in WT mice (Fig. 3F-3I). Two types of abnormal Pac (Pac-like 1 and Pac-like 2) were observed in KO mice. Although the discontinuous SYCP3 signal was observed in most chromosomes, it was much more fragmented in Pac-like 2 cells (Fig. 3G). Moreover, SYCP3 signal in the form of bubbles and forks, which indicate large scale asynapsis between homologous chromosomes, was also frequently observed in Pac-like 2 (Fig. 3G). Discontinuous SYCP3 signals were more readily detected in chromosomes in KO Dip (Fig. 3H). Averagely, 33.5% of Pac (10.5% Pac-like 1 and 23.0% Pac-like 2) and 55% of Dip in KO mice contained abnormal SYCP3 staining signals; while, the percentage was rather low in WT littermates (about 0.6%) (Fig. 3I). Moreover, the percentages of different SCs were changed in KO mice with Lep/Zyg enrichment and Pac/Dip depletion (Fig. 3J). Interestingly, γH2AX signals all disappeared in autosomes and still existed in XY body in late Pac and Dip from KO mice, suggesting that DSB repair was not affected by *miR-202* KO (Fig. 3G, H). These results indicated that *miR-202* plays an important role during meiosis I prophase by regulating synapses of homologous chromosomes.

### Loss of *miR-202* results in inter-sister synapsis and reduced crossover formation

Asynapsis in KO Pac was confirmed by the co-immunostaining of SYCP3 and SYCP1 (Fig. 4A). In Pac, X and Y chromosomes undergo partial synapsis in the pseudoautosomal region (PAR). Interestingly, co-localization of SYCP3 and SYCP1 in X and Y chromosomes in many KO Pac was also detected beyond PAR. The percentage of such cells in KO mice was 77.6% (n = 98 nuclei, Fig. 4B) while that in WT mice was only 4.3% (n = 46 nuclei). A closer examination of the autosomal bivalents revealed that, in some Pac, SYCP3 and SYCP1 co-localization was also detected in the separated axis of the bubble and fork regions (Fig. 4A). Co-localization of SYCP3 and SYCP1 in asynapsed homologs is a sign of inter-sister chromatid synapses and has been reported in several mutant mouse lines with total or partial removal of certain cohesin proteins (5, 7-12). The percentage of these cells was 19.4% in our KO mice (n = 98 nuclei, Fig. 4B) and 0% in WT ones (n = 46 nuclei). This means that 84.3% (19.4% divided by 23.0%) of Pac-like 2 cells underwent inter-sister synapsis in their autosomes. Moreover, 31.6% of Pac, in which inter-sister synapsis was detected in sex chromosomes, also underwent inter-sister synapsis in the asynapsed regions of the autosomal homologous chromosomes (n = 76 nuclei). X and Y chromosome inter-sister synapsis was more evident when super-resolution structured illumination microscopy (SIM) was used, as the tripartite structure of the synaptonemal complex on X and Y chromosomes beyond PAR was observed in KO Pac (Fig. S5A). The co-localization of SYCP3 and SYCP1 on asynapsed chromosomes was observed as early as in KO Zyg (Fig. S5B).

**Fig. 4.**
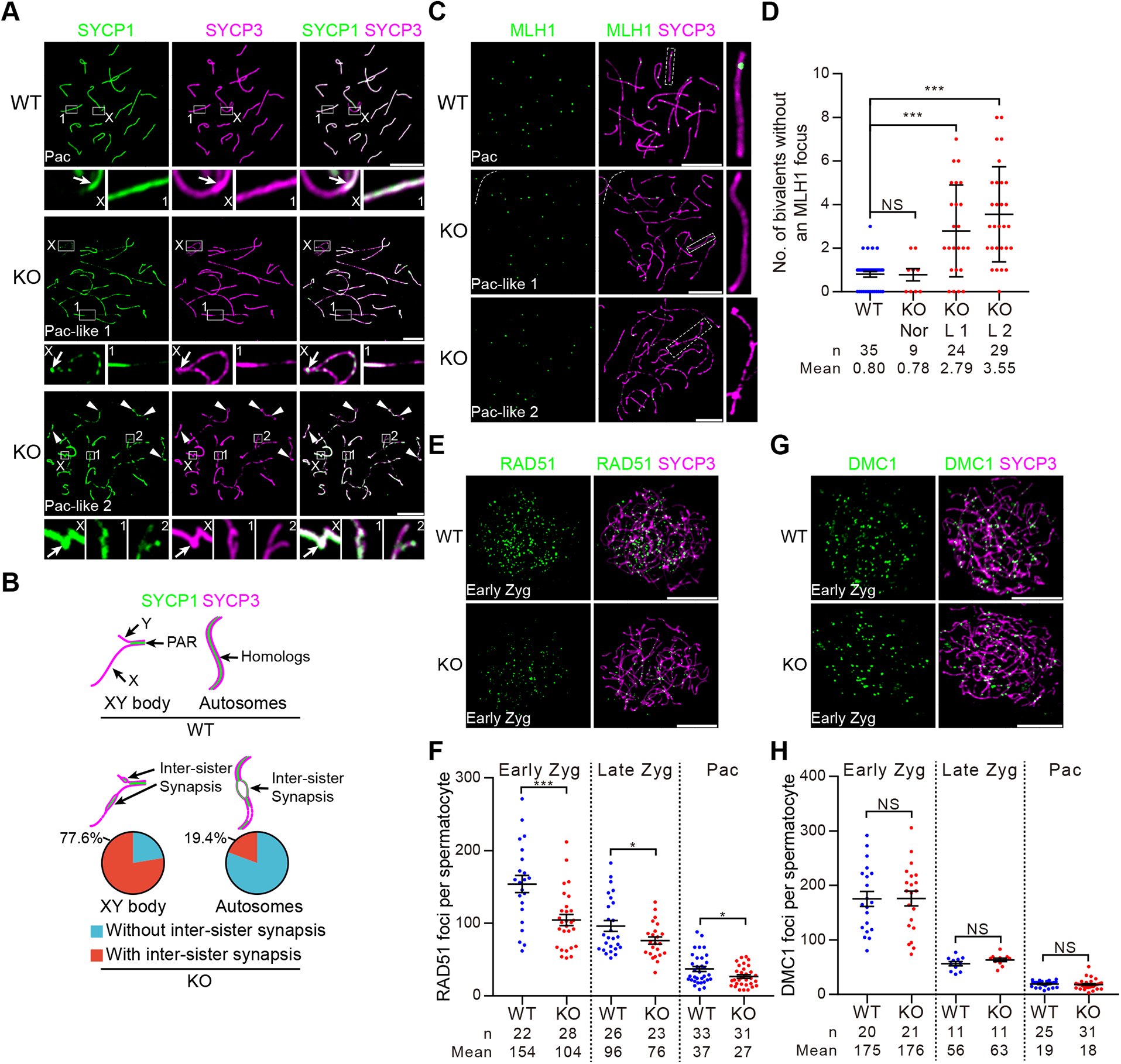
Loss of *miR-202* causes inter-sister synapsis and impaired crossover formation. (**A**) Nuclear-spread SCs from WT and KO mice were immunostained for SYCP3 and SYCP1. Representative images of WT and KO Pac are shown. Enlargements of boxes are shown below the respective full nucleus images, where “X” marks the XY body regions. Arrows indicate the PAR. Arrowheads indicate inter-sister synapsis besides the boxed regions. (**B**) Schematic representation shows the SYCP3 and SYCP1 patterns for XY body and autosomes in WT and KO Pac in (A). In the lower panel, percentages of KO pachynema with and without inter-sister synapsis in XY body and autosomes are respectively shown. At least three mice of each group were counted. (**C**) Nuclear-spread SCs from WT and KO mice were immunostained for SYCP3 and MLH1. Representative images of Pac are shown. Magnified views are indicated by dashed areas. (**D**) Number of bivalents without an MLH1 focus per Pac in (C). At least three mice of each group were counted. Nor, normal Pac; L 1, Pac-like 1; L 2, Pac-like 2. (**E, G**) Nuclear-spread spermatocytes from WT and KO mice were immunostained for SYCP3 together with RAD51 (E) or DMC1 (G). Representative images of early zygonema are shown. (**F, H**) Quantification of RAD51 (F) or DMC1 (H) focus numbers of early Zyg, late Zyg, and Pac. At least three mice of each group were counted. All values are shown as mean ± SEM. *p < 0.05, ***p < 0.001. NS, not significant. Scale bars: 10 μm.

As defective synapsis usually causes abnormal homologous recombination (4), we next tested whether this is the case upon *miR-202* KO by examining the immunostaining signals of several marker proteins. MLH1 marks crossovers formed between homologous chromosomes in Pac as a result of homologous recombination, and 1∼2 MLH1 foci are detected on each bivalent (7, 42). Indeed, there were consistently more chromosomal bivalents lacking MLH1 foci in KO cells of Pac-like 1 and Pac-like 2 than in the WT ones (Fig. 4C and 4D). In addition, the average number of MLH1 foci per WT Pac was 22.1. In contrast, the number dropped to 20.1 and 19.6 in Pac-like1 and Pac-like 2, respectively (Fig. S5C). RAD51 and DMC1 are associated with the single-stranded 3’ ends of the resected DSBs and can catalyze strand exchange in Lep/Zyg (43, 44). For early and late Zyg, and Pac, the numbers of the RAD51 foci were all significantly lower in the KO mice than in the WT mice (Fig. 4E and 4F). Interestingly, the numbers of the DMC1 foci in these cell types were similar between WT and KO mice (Fig. 4G and 4H). These results demonstrate that *miR-202* ensures synapsis and recombination between homologous chromosomes by preventing inter-sister synapsis.

### REC8 is progressively lost due to elevated expression of separase in *miR-202* KO mice

As the phenotypes of *miR-202* KO mice are similar to those of KO mice of the cohesin proteins (5, 7-12), we next examined whether the localization and expression of meiosis-specific cohesin proteins, including REC8, RAD21L, SMC1β and STAG3, is affected by *miR-202* KO. We selected REC8 for the following reasons: 1) In *Rec8* KO SCs, the entire chromosome axis becomes two separate SYCP3-labeled structures, despite the fully localized RAD21L, while the chromosome axis is regionally separated in hypomorphic *Stag3* mutant and *Smc1β* KO spermatocytes, in which REC8 is partly reduced (9, 10); 2) *Rad21l* KO SCs display impaired synapsis, where SCs form predominantly between non-homologous chromosomes (11, 12). Consistent with previous studies (9, 10), REC8 signals appeared as foci that co-localized with SYCP3 from Lep to Dip in WT mice (Fig. 5A-5D). A similar localization was observed in KO mice. However, REC8 staining was weaker in KO Zyg/Pac/Dip than in WT ones, particularly on the REC8 and SYCP3 co-staining images under high-magnification (Fig. 5A-5D). For quantitative comparisons, we measured the inter-focus distances of REC8 signals instead of the number of foci because foci often clustered together and were hard to count (9, 10). We found that the average distance of REC8 signals in KO mice was not significantly different from that in the WT mice in Lep. However, the difference was significant in Zyg and Pac, and was the most significant in Dip (Fig. 5E). Therefore, it appeared that REC8 underwent progressive loss during the prophase of meiosis I. Moreover, Western blot analyses of FACS-isolated SCs showed that REC8 decreased significantly in the KO mice (Fig. 5F-5G and S6A-S6C). These results reveal that the inter-sister synapsis in *miR-202* KO mice might be a consequence of REC8 loss.

**Fig. 5.**
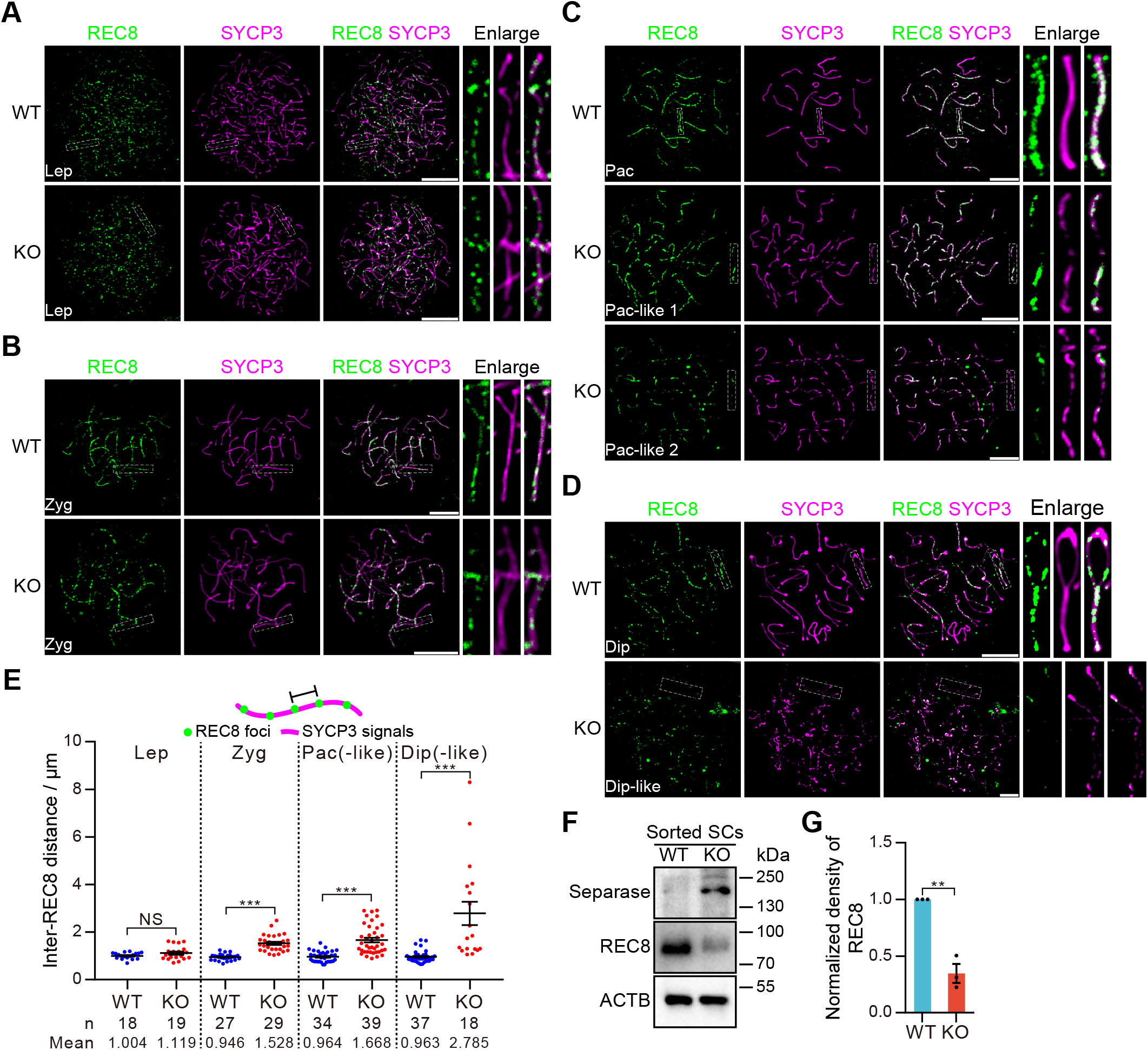
REC8 is progressively lost in *miR-202* KO mice. (**A**-**D**) Representative nuclear spreads of WT and KO spermatocytes at Lep (A), Zyg (B), Pac (C) and Dip (D). SCs were immunostained for REC8 and SYCP3. REC8 foci distribute along SYCP3-labelled structures. Magnified views are indicated by dashed areas. (**E**) Inter-REC8 distances along axial elements. Five stretches or homologs were measured and the average distance was calculated for each nucleus. Each point represents one nucleus. The upper panel is the schematic representation for distance measurement. At least three mice of each group were counted. (**F**) Western blot analyses for REC8 and separase in spermatocytes isolated by FACS after Hoechst staining. n = 3. (**G**) Normalized density of REC8 in (F). All values are shown as mean ± SEM. **p < 0.01, ***p < 0.001. NS, not significant. Scale bars: 10 μm.

### Separase-mediated cleavage of REC8 is aberrantly activated in *miR-202* KO mice

Under normal condition, REC8 on chromosome arms is cleaved by separase at the metaphase-anaphase transition of meiosis I, so that homologous chromosomes originally maintained as pairs by the crossovers can separate from each other (13, 14, 45). The proteomic data (ProteomeXchange, PXD005552) in our previous study showed that the protein level of separase in cultured spermatogonial stem cells (SSCs) increased significantly upon *miR-202* KO (27). Therefore, we next examined if the localization and/or amount of separase would change in *miR-202* KO mice. No immunostaining of separase was detected on the chromosome spreads of any WT prophase SCs (Fig. 6A-6D). Surprisingly, the high level of separase signal was detected in the chromosome spreads of the KO SCs from Zyg to Pac (Fig. 6B-6D). Separase partially co-localized with SYCP3 in the KO mice (Fig. 6B-6D), and this is consistent with the previous finding that cohesin cleavage by separase requires DNA in a sequence-nonspecific manner (46). Moreover, western blot analyses of FACS-isolated SCs confirmed that separase increased dramatically in the KO mice (Fig. 5F-5G and S6A-S6C).

**Fig. 6.**
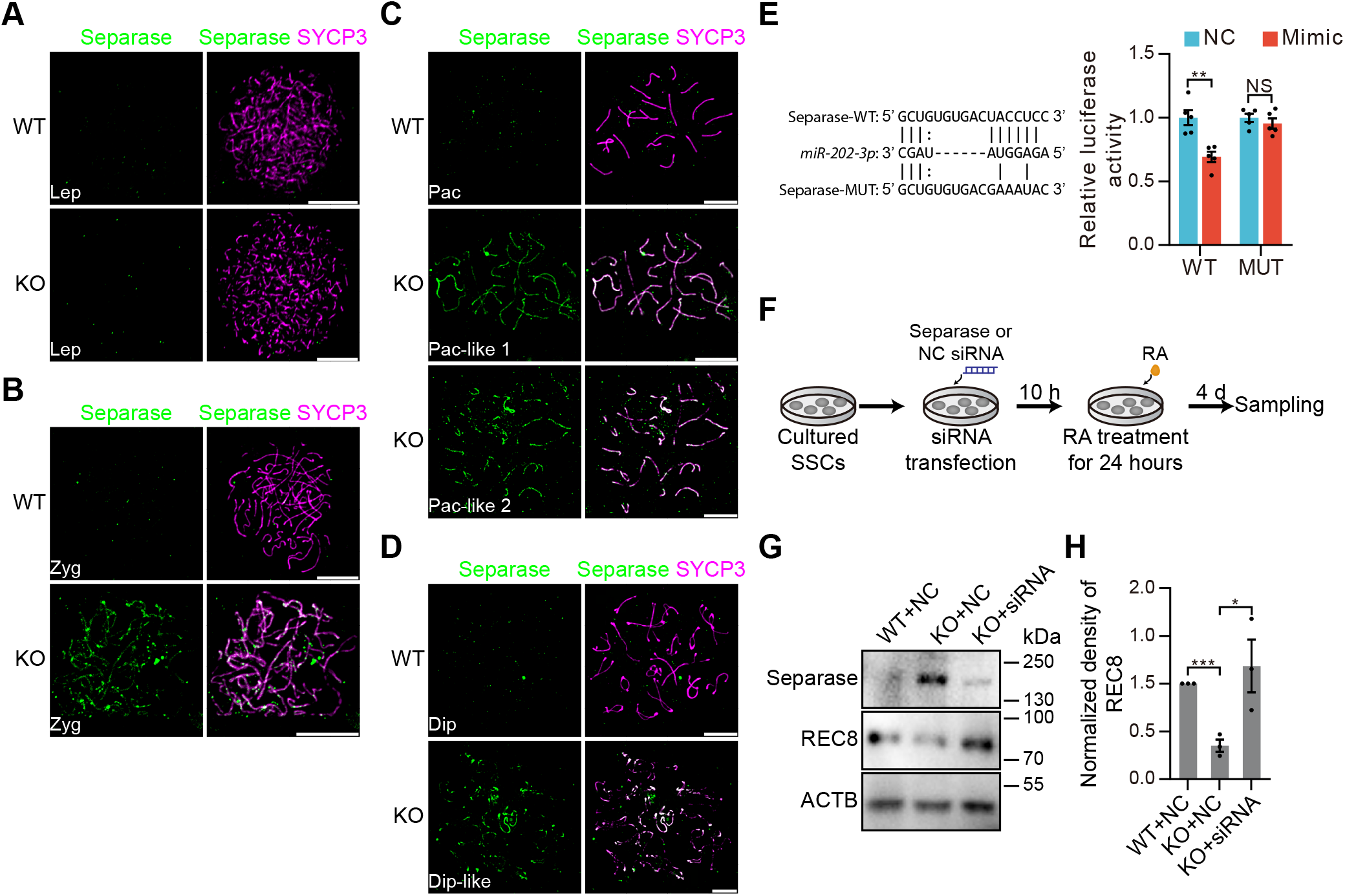
Separase-mediated cleavage of REC8 is aberrantly activated in *miR-202* KO mice. (**A**-**D**) Representative nuclear spreads of WT and KO SCs at Lep (A), Zyg (B), Pac (C) and Dip (D). SCs were immunostained for separase and SYCP3. Separase signals distribute along SYCP3-labelled structures from Zyg to Dip in KO. (**E**) Validation of separase mRNA as a direct target of *miR-202-3p* by dual luciferase assay. Data are normalized to the scrambled negative control (NC) mimic. Left panel: the sequence of *miR-202-3p*, and predicted miRNA regulatory elements at the 3’ UTR of separase mRNA and mutated 3’ UTR sequence. (**F**) Diagram for verification of separase activation and REC8 cleavage in cultured SSCs. SSCs were transfected with separase or NC siRNA. Ten hours later, the cells were treated with RA (retinoic acid) for 24 hours. Four days after induction, the cells were harvested and analyzed. (**G**) Western blot analyses for REC8 and separase for cells in (F). n = 3. (**H**) Normalized density of REC8 in (G). All values are shown as mean ± SEM. *p < 0.05, **p < 0.01, ***p < 0.001. NS, not significant. Scale bars: 10 μm.

Interestingly, separase mRNA was found to be a predicted target of *miR-202-3p* (Fig. 6E) (47). Dual luciferase assay revealed that the 3’ UTR of separase that contained the predicted target site was actually targeted by the *miR-202-3p* mimic. As a negative control, the mutated target sequence was not targeted by *miR-202-3p*, indicating that separase mRNA was a direct target of *miR-202-3p* (Fig. 6E). We previously reported that long-term cultured SSCs could be induced to Lep/Zyg upon retinoic acid treatment (48). We then used this *in vitro* system to further test whether REC8 degradation is mediated by separase (Fig. 6F and S6D). Consistent with the western blot results using the isolated SCs, REC8 and separase exhibited reciprocal changes in induced KO cells compared with induced WT ones. Importantly, both changes in KO SSCs were reverted by the pretreatment of SSCs with a siRNA specific to the separase mRNA but not with a nonspecific randomly scrambled small RNA (Fig. 6G-6H and S6E). Taken together, *miR-202* KO results in the elevated expression of separase, which in turn leads to precocious degradation of REC8 during the prophase of meiosis I.

## Discussion

Meiosis is a key step of gametogenesis and involves many important processes such as synapsis and DNA recombination that are regulated by a multitude of proteins (1-3). Cohesin plays important roles in synapsis, DNA recombination, and segregation of homologous chromosomes (5, 7-12). In the present study, we found that the KO of *miR-202*, a miRNA that is evolutionarily conserved and highly expressed in the testis, resulted in reduced male fertility, reduced numbers of meiotic and post-meiotic germ cells, and many molecular defects including inter-sister synapsis and impaired crossover formation during meiosis prophase I. More importantly, we revealed that loss of *miR-202* resulted in elevation of separase and reduction of REC8 and that separase mRNA was a direct target of *miR-202-3p* (Fig. 7). These results provide new insights into the regulatory network of meiosis in which miRNAs play important roles.

**Fig. 7.**
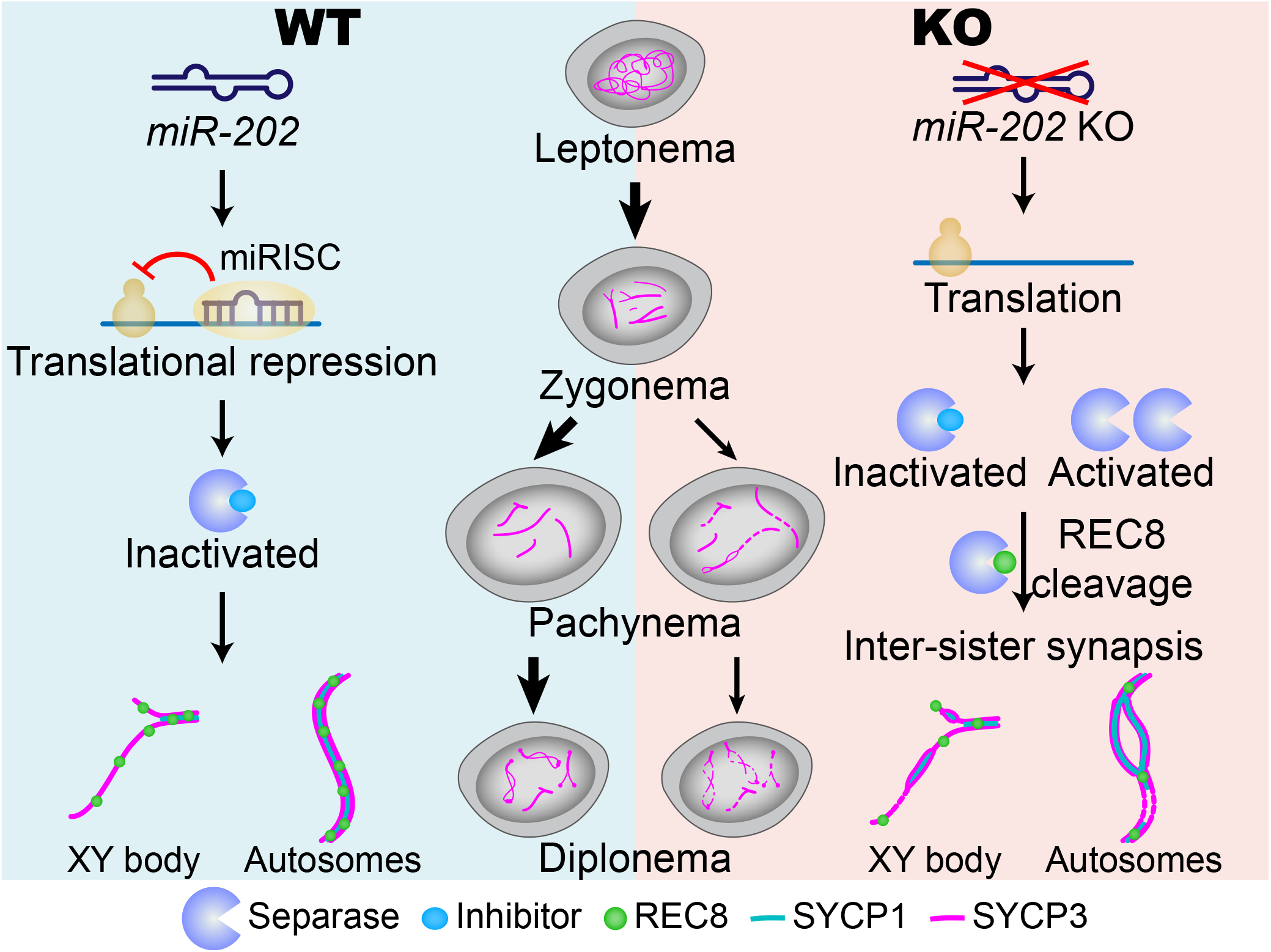
*miR-202* safeguards meiosis prophase I by preventing premature degradation of REC8 mediated by separase. During meiosis prophase I, the activity of a protease, separase, is strictly inhibited by its inhibitory partners such as securin and cyclin B, and *miR-202*-mediated repressed translation via direct target of separase mRNA, to safeguard the REC8 level for homologous synapsis and recombination. Loss of *miR-202* elevates the level of separase, which outnumbers its inhibitors and cleavages its meiotic substrate REC8 prematurely, causing partial arrest of meiosis at the Zyg/Pac transition and inter-sister synapsis.

The opinion that miRNAs are important for mammalian spermatogenesis is mainly based on three types of observations: expression dynamics in testes and spermatogenic cells (22), observations made using *in vitro* cultures of spermatogonia (27, 49, 50), and phenotypic evaluations of KO mice of genes such as *Dicer, Drosha*, and *Dgcr8* that encode key enzymes/regulators in miRNA biogenesis (17-25). Few mouse KO models of miRNA functions in spermatogenesis have been reported so far and they are all for miRNA gene clusters but not for single miRNAs. For example, the conditional KO of the *miR-17-92* cluster, which contains 6 miRNA genes (*miR-17, 18a, 19a, 20a, 19b-1, 92a-1*), in spermatogenic cells, resulted in smaller testes, lower sperm concentration, and many degenerating tubules containing only Sertoli cells. Interestingly, the litter size was not significantly different from that of the WT mice (51). The *miR-34* family includes 6 miRNAs (*miR-34a, b, c* and *449a, b, c*) that are generated from three loci: *miR-34a, miR-34b/c* and *miR-449*. Surprisingly, all the KO mice of the three single loci, and those KO mice of the *miR-34a* and *miR-34b/c* double loci exhibited normal spermatogenesis and fertility (52, 53). Oligoasthenoteratozoospermia and infertility was only observed in the mice with *miR-449* and *miR-34b/c* double loci KO (54-56). Moreover, the KO of the *miR-379*/*miR-544* cluster (∼9.4 kbp) in spermatogenic cells did not generate any observable phenotype (57).

We previously reported that 43% of small RNA reads in type A spermatogonia was contributed by miRNAs, but this number dropped to 7% and 5% in Pac and rSTs, respectively (58). In vertebrates, *miR-202* is a member of the *let-7* family (26), and is located in the intergenic region as a singleton in the genomes of mice and humans. Due to its membership in the well-studied *let-7* family and its isolated genomic location, we selected *miR-202* for functional studies. By using an inducible CRISPR-Cas9 system established in mouse SSC lines, and the transcriptomic and proteomic methods, we previously showed that *miR-202* played an important role in maintaining the undifferentiated state of cultured SSCs by targeting diverse genes post-transcriptionally (27). In the present study, we studied the *in vivo* function of *miR-202* by evaluating the phenotypic and molecular changes of its KO mice and found it played an important role specifically in spermatogenesis. Therefore, these findings may contribute to the understanding of the functions of single miRNA genes and the mechanism of meiosis.

Meiosis consists of inter-dependent processes that must be intricately coordinated by a large number of regulators (1-3). For example, synapsis and recombination of homologous chromosomes are tightly coupled by protein complexes such as synaptonemal complex, cohesins, and recombinases, which interact with each other by complicated mechanisms yet to be fully understood. The disruption of one process usually affects the others. Therefore, genetic ablations/mutations of most regulator genes usually share common abnormalities in synapsis, DNA DSB repair, and crossover formation, accompanied by the deaths of meiotic cells (1-3). In this sense, it is not surprising that the *miR-202* KO mice in this study also displayed multiple defects in these processes (Fig. 3 and 4). On the other hand, as meiosis progresses in a step-wise manner and different regulators may exert their functions at one or more specific steps, several meiotic arrest stages have been revealed by using mouse genetic models (1). The Zyg/Pac transition is the most frequently reported arrest stage probably because DSB repair, synapsis, and crossover formation all have to be completed in Pac; otherwise, either Pac cannot be generated or defective ones are eliminated by apoptosis (1). Such arrest may not be a clear cut probably and mainly because of redundancy and complimentary effects of family members of the regulators. Therefore, it is also not surprising that meiosis in *miR-202* KO mice of this study arrested at Zyg/Pac transition incompletely but significantly as evidenced by changes in cellular composition and molecular marker localizations (Fig. 2F, 3J, 4A-4B and S5A-S5B).

Two observations related to DNA DSB repair and recombination were noteworthy. First, although the number of Zyg marked by the γH2Ax diffuse staining was increased significantly in KO mice (Fig. 2F and 3J), DSB repair in autosomes seemed to be well accomplished in Pac-like cells as γH2Ax signal was only seen in the sex body (Fig. 3G and 3H) despite the detection of asynapsis in both sex chromosomes and autosomes (Fig. 4A and 4B). Second, the numbers of MLH1 and RAD51 foci but not those of the DMC1 foci were reduced by *miR-202* KO in Pac-like cells (Fig. 4C-4H). This observation was supported by two previous studies (44, 59). One demonstrated that KO of both *Rad21l* and *Rec8* reduced RAD51 but not DMC1 (59). Another one showed that DMC1, not RAD51, performed strand exchange during DSB repair (44). Taken together, it seems that *miR-202* plays a more important role in synapsis than in DSB repair, which was only delayed but not totally blocked during the Zyg/Pac transition in the KO mice. Alternatively, DSBs are not repaired in Pac-like cells but cannot be observed because such cells are quickly eliminated by apoptosis as the apoptotic cells were Pac in the KO mice (Fig. 1K-1M).

Among all the molecular defects in the meiocytes of *miR-202* KO mice, the inter-sister synapsis is of particular interest as it was previously only observed in mutant mice of cohesin protein genes (5, 7-12). This leads us to investigate the relationship between *miR-202* and REC8, of which the KO mice were first reported to undergo inter-sister synapsis (7, 8). This exploration turned out to be fruitful with several unexpected findings: 1) We found that the number of REC8 foci (indicated by the inter-foci distance in Fig. 5E as this measurement was easier to perform) in chromosomes and the total REC8 level were both reduced significantly in *miR-202* KO mice (Fig. 5F-5G) while these two parameters of separase were changed significantly in the opposite direction (Fig. 5F-5G, and 6A-6D); 2) Separase mRNA was a direct target of *miR-202-3p* (Fig. 6E); 3) The total protein changes of both REC8 and separase were partially rescued by the siRNAs that targeted separase mRNA in induced SCs (Fig. 6F-6H). The degradation of REC8 at the end of meiosis I prophase is mediated by separase, which is activated by the ubiquitin-dependent degradation of its inhibitory partners, such as securin and cyclin B (14, 45, 60). Pre-activation of separase due to overexpression has been reported in cancer cells and oocytes, which induces aneuploidy (61-63). It is highly likely that elevated level of separase in *miR-202* KO SCs outnumbers its inhibitors and cleavages its meiotic substrate REC8 prematurely in meiosis prophase I (Fig. 7).

Taken together, our findings indicate that *miR-202* is a novel regulator of mammalian meiosis that directly interacts with the separase-REC8 axis, which is key to synapsis and recombination during meiosis. As miRNAs usually have multiple targets, the mechanism underlying the regulatory roles of *miR-202* may be better understood by identifying more targets. Moreover, it is highly likely that *miR-202* acts at multiple stages of spermatogenesis as suggested by our previous *in vitro* study (27). Therefore, it will be important to examine whether stages of spermatogenesis other than meiosis are disrupted in the *miR-202* KO mice.

## Materials and Methods

### Mice

All of the animals used in this study were approved by the Animal Ethics Committee of the Institute of Zoology at the Chinese Academy of Science. All of the procedures were conducted in accordance with institutional guidelines. Animals were specific-pathogen free (SPF). All mice had access to food and water ad libitum, were maintained on a 12:12 hours light-dark artificial lighting cycle, with lights off at 19:00, and were housed in cages at a temperature of 22-24°C.

Mice were generated through the CRISPR/Cas9 gene-editing approach (64). The gRNAs (gRNA-1s: TTGCAGGGGATCAATCCTTC<u>TGG</u>; gRNA-1a: GCTAATGACTTTGTTTGGGT<u>GGG</u>; PAM sites of sgRNAs are underlined) that were designed in our previous study (27). The double-stranded DNA of sgRNAs annealed from synthesized complimentary oligonucleotides were inserted into the pUC57-sgRNA vector (Addgene, 51132), and gRNAs were transcribed *in vitro*. Cas9 mRNA was transcribed *in vitro* from pST1374-NLS-flag-linker-Cas9 (Addgene, 44758). Then, Cas9 mRNA and sgRNAs were injected into zygotes from mating C57BL/6J males with superovulated C57BL/6J females. We obtained two male founders (F0) identified by genomic PCR (Table S3). We then crossed both males with females and the one that transferred the KO allele to the offspring were used for further study.

All of the mice were maintained on a C57BL/6J;ICR mixed background (Fig. S1A). The deletion of the *miR-202* cassette was verified by genomic PCR and qPCR (Fig. S1B-C).

### Cell clustering and re-clustering of scRNA-seq data

The filtered gene expression data were analyzed by the Seurat package (Version 3.2.2) (65). Global scaling normalization was applied with the “NormalizeData” function via the “LogNormalize” method with a scale factor 10,000. The top 2000 variable genes were identified by the “FindVariableGenes” function and used for principal component analysis. Significant principal components that were enriched with low p value (<1×10^−5^) genes were used for cell clustering. Using the graph-based clustering approach implemented in the “FindClusters” function of the Seurat package, with a resolution of 0.2, a seed of 323 and otherwise default parameters, cells were clustered and re-clustered by UMAP aligned coordinates and the resulting clusters were consistent with the mapping of known markers (29).

### Antibodies

All antibodies used in this study are listed in Table S4. We obtained the and the anti-H1t antibody from Prof. Mary Ann Handel in Jackson Laboratory (41), as a generous gift.

### Immunostaining of spermatocyte chromosome spreads

Spermatocyte spreads of the testicular samples were performed using the drying-down technique previously described by Peters *et al*. (66). Briefly, the testes were dissected, and the seminiferous tubules were washed in PBS. Then, tubules were placed in a hypotonic extraction buffer containing 30 mM Tris, 50 mM sucrose, 17 mM trisodium citrate dihydrate, 5 mM EDTA, 0.5 mM DTT, and 0.5 mM PMSF (pH 8.2), for 30-60 min. Subsequently, the tubules were torn to pieces in 0.1 M sucrose (pH 8.2) on a clean glass slide and were pipetted repeatedly to make a suspension. The cell suspensions were then dropped on slides containing 1% PFA and 0.15% Triton X-100 (pH 9.2). The slides were dried for at least two hours in a closed box with high humidity. Finally, the slides were washed twice with 0.4% Photo-Flo 200 (Kodak) and dried at room temperature. The dried slides were stored at −20°C for the immunofluorescent staining. Slides of meiotic chromosome spreads were performed using the procedures previously described (67). The rabbit anti-SYCP3 antibody was used only when co-staining of SYCP3 and MLH1 was performed.

Immunolabeled chromosome spread nuclei were imaged on confocal laser scanning microscopes (Leica or Carl Zeiss) using 63× oil-immersion objective, unless otherwise stated. Structured illumination microscopy (3D-SIM) was performed on SIM super-resolution microscopes (Nikon or Carl Zeiss) 100× oil-immersion objective.

### Dual luciferase assay

We used microT-CDS to predict mRNA target sites of *miR-202* (47). The predicted 3’ UTR for separase was amplified from mouse testis cDNA (Table S3) and inserted into the pMIR-REPORT Luciferase vector.

Furthermore, dual luciferase assays were conducted following our previous report (27). Briefly, the 293FT cells were plated in 96-well plate pretreated with 0.2% gelatin. About 24 hours later, the *miR-202* or scrambled NC mimics of 100 nM were first transfected using the Lipofectamine RNAiMAX Reagent (Invitrogen). Ten hours later, 50 ng of the Firefly Luciferase report plasmids (pMIR-REPORT Luciferase) and 5 ng of the Renilla Luciferase internal control plasmid (pRT-TK) were co-transfected by the Lipofectamine 2000 Reagent (Invitrogen). Fourty-eight hours after plasmid transfection, luciferase activity was examined using the Dual-Luciferase Report Assay System (Promega) on the Synergy4 (Bio-Tek) platform. Data were first normalized to empty the vector and then to mimic the negative control.

### Statistical analysis

All experiments reported here were repeated at least three independent times. All of the values in the figures are shown as mean ± SEM unless otherwise stated. Excel 2016 or GraphPad Prism 7 was used to perform statistical analyses. For statistical analysis of differences between two groups, two-tailed unpaired Student’s t tests were used. For statistical analysis of focus number comparisons for MLH1, RAD51 and MLH1, nonparametric one-tailed Mann-Whitney tests were used. For statistical analysis in Fig. 2C and 2F, Chi-square test was used. For statistical analysis in Fig. 2H, Wilcoxon rank-sum test was used. No samples or animals were excluded from analyses. Sample size estimates were not used. Mice analyzed were litter mates and sex-matched whenever possible. Investigators were not blinded to mouse genotypes or cell genotypes during experiments.

In all figures, *, ** and *** represent p < 0.05, p < 0.01, and p < 0.001, respectively. NS (not significant) indicates not statistically significant difference (p > 0.05).

## Data availability

The single-cell RNA-seq data from this study will be submitted.

## Acknowledgments

We thank Mary Ann Handel for the H1t antibody. We thank Bing Zhou, Maria Jasin and Nicolas Plachta for their critical suggestions. We thank Shiwen Li, Shuguang Duo, Xili Zhu, Xia Yang, Hua Qin and Qing Meng in Institute of Zoology, Chinese Academy of Sciences for their technical assistance. We thank LetPub (www.letpub.com) for its linguistic assistance during the preparation of this manuscript.

This work was supported by grants from the Ministry of Science and Technology of China (2016YFC1000606, 2018YFE0201100, 2015CB943002), and the National Natural Science Foundation of China (31771631, 31970795).

## Conflict of Interest

No competing interests declared.

## Author Contributions

C.H. supervised the project, conceived and designed the study, analyzed the data and wrote the manuscript. J.C. conceived and designed the study, performed most experiments, analyzed the data and wrote the manuscript. C.G. and X.L. conducted bioinformatic analyses. C.Z., Y.N., L.M., W.H. and D.X. performed experiments. K.L. revised the manuscript. All authors commented on the manuscript.

## Notes

### Competing Interest Statement

The authors have declared no competing interest.

