## Supplemental Information for "MicroRNA-202 safeguards meiotic progression by preventing premature degradation of REC8 mediated by separase"

### Supplementary Information

#### Supplementary Materials and Methods

##### Fertility test

*MiR-202* WT or KO adult male mice were each housed with at least three eight-week-old C57BL/6J WT female mice for one month. Female mice were checked for the presence of a vaginal plug every morning to examine mating activity. The female mice that were vaginal-plug positive were separated and observed for three weeks for pregnancies. The pregnancy rate and the number of litters per female were recorded and calculated.

##### Sperm count

The cauda epididymis was dissected and immediately minced in 1 ml of PBS solution. Semen was then released (15 minutes at 37ºC, 5% $CO_2$) from three to five small incisions. Thereafter, the total sperm count was assessed with a hemocytometer.

##### Histological analyses

Testes dissected from the wildtype and KO mice immediately after euthanasia were fixed in 4% paraformaldehyde or Bouin's Solution for up to 24 hours, dehydrated using gradient concentration of ethanol, treated with xylene, and then embedded in paraffin. Five-micrometer-thick sections were cut and mounted on glass slides. After deparaffinization in xylene and re-hydration in gradient concentration of ethanol, the Bouin-fixed testis cross-sections were used for Hemotoxylin and Eosin (HE) staining, and PFA-fixed sections were used for immunohistochemistry (IHC), immunofluorescence (IF) analyses, and TUNEL assays.

##### Generation of cell suspensions

Single-cell suspensions were generated from adult WT and KO mouse testes using a two-step enzymatic digestion approach (1). Mice were sacrificed, and then the testes were removed and decapsulated. The seminiferous tubules were cut into small pieces and incubated in DPBS containing 1 mg/ml Collagenase I (Gibco) at 37ºC for five minutes with gentle shaking. The dispersed seminiferous tubules and cells were collected by centrifugation at 100 g for two minutes at 4ºC. The pellet was washed once with DPBS and digested in 0.25% Trypsin (Gibco) containing 1 mg/ml DNase I (AppliChem) at 37ºC for five minutes with gentle shaking. The suspension was added with FBS (Vistech) to terminate digestion. Then, cells were collected by centrifugation at 600 g for five minutes and washed with 0.04% BSA/DPBS. The cells were then filtered through a 40-μm cell strainer (BD Falcon), and cell debris was removed by FACS. The cells were resuspended at a concentration of 1 million cells per ml in 0.04% BSA/DPBS.

**10× Genomics single-cell RNA-Seq**

Chromium Next GEM Single-cell 3' GEM Library and Gel Bead Kit V3.1 (10× Genomics, 1000121) and Chromium Next GEM Chip G Single Cell Kit (10× Genomics, 1000120) were used for cell capture and cDNA synthesis. The cell suspension was loaded onto the Chromium single-cell controller (10× Genomics) to generate single-cell gelbead emulsions (GEMs) according to the manufacturer's instruction (2). About 20,000 cells were added to each channel, and the target recovery rate was estimated to be 13,000 cells. Captured cells were lysed and the released RNA was barcoded through reverse transcription in individual GEMs. Reverse transcription was performed using a T series Multi-Block Thermal Cycler (LongGene). The following procedure was used: 53 °C for 45 min, 85 °C for 5 min, and 4 °C forever. The quality of the generated cDNA was assessed by the Agilent 2100, which was performed by Emei Tongde (Beijing, China). Sequencing (performed by NovoGene, Beijing) was performed on the Illumina NovaSeq 6000 sequencer with 150 bp (PE150) paired-end reads.

**Processing raw data from scRNA-seq of 10× Genomics**

Trimmed FASTQ files (26 bp cell barcode and UMI Read1, 8 bp i7 index, and 100 bp Read2), were generated using the CellRanger (Version 1.3.1) "mkfastq" command. The Cell Ranger "mkref" function with default settings was used to process the genomic sequence (Mus musculus genome GRCm38) and the annotation file for read alignment. We removed cells with low quality using the following thresholds: 1) UMIs or gene numbers beyond 3 median absolute deviations away from the median determined by the "isOutlier" function of the Scater package (Version 1.14.6); 2) the ratio of mitochondria over 0.2.

**Identification of differentially expressed genes (DEGs)**

DEGs between WT and KO spermatocytes from scRNA-seq were identified by the "FindMarkers" function of the Seurat package using the "Wilcox" test, which returned "p_val_adj" using the Bonferroni correction and the log-transformed fold change, "avg_logFC", and the cutoff was set to p_val_adj < 0.05.

**GO analysis**

Gene ontology (GO) analysis was performed using DAVID bioinformatics tools (3) and GO terms with $p<0.05$ were considered to be significant.

**Section immunostaining and TUNEL assay**

Deparaffinized sections were boiled for 15 min in a sodium citrate buffer for antigen retrieval. Then, the slides were incubated with primary antibodies overnight at 4ºC and then incubated with secondary antibodies for two hours at room temperature. For IF, signals were visualized by conjugating fluorophore with the secondary antibodies. For IHC, the sections were stained with a horseradish peroxidase (HRP)-conjugated secondary antibody. We examined the SYCP3 expression patterns using the mouse anti-SYCP3 antibody. Nuclei were counterstained with DAPI and hematoxylin for IF and IHC, respectively. After staining, the IHC and IF sections were examined with a Nikon microscopy and confocal laser scanning microscopes (Leica or Carl Zeiss), respectively. The TUNEL assay was conducted using the DeadEnd Fluorometric TUNEL System (Promega) according to the standard protocol.

**Stage characterization for nuclear-spread spermatocytes**

Leptonema are characterized by the presence of small stretches of SYCP3. At early zygonema, homolog synapsis initiates and longer SYCP3 stretches are visible due to the continuous elongation of chromosome axes. At late zygonema, chromosome axis formation completes at the same time that more than 50% homologs synapsis. At pachynema, homologs are fully synapsed, except in the non-pseudoautosomal region of the XY chromosomes. Because of chromatin condensation, pachytene chromosome axes are thicker and shorter. At diplonema, chromosome desynapsis ensues. At this stage, homologs are held together by the recombination sites (crossovers), which are recognized as chiasmata (4). Images were processed using Image J to count focus numbers and only foci colocalizing with the chromosome axes were counted. The inter-REC8 distances were measured by manually tracing the chromosome axes in FIJI (5).

**Western blotting**

Spermatocytes sorted by Hoechst 33342 (Sigma-Aldrich) staining were directly lysed in a RIPA buffer supplemented with Protease Inhibitor Cocktail. Lysates were then centrifuged at 20,000 g for 10 minutes at 4°C, and supernatants were used for Western blot analyses. Briefly, lysates were run in SDS-PAGE gel and transferred to PVDF membranes. The blots were blocked with 5% BSA for 2 hours, incubated with primary antibodies overnight at 4ºC and then incubated with HRP-conjugated secondary antibodies at room temperature for 2 hours. The proteins were detected using SuperSignal West Pico PLUS Chemiluminescent Substrate (Thermo Fisher) on the ChemiDoc XRS$^{+}$ system (Bio-Rad). The band densities were analyzed by ImageJ.

**Isolation of SCs**

We isolated SCs from the testes of four-month-old mice. Single-cell suspensions were generated as described above. After filtered, the cells were incubated in DMEM containing 1 μg/ml of Hoechst 33342 (Sigma-Aldrich) at 37ºC for 30 minutes. The cells were sorted by a MoFol XDP instrument (Beckman Coulter). The cells were first sorted to collect tetraploid cells and were further sorted into two population based on their forward scatter (FSC) and side scatter (SSC) features. The population of larger sizes consisted mainly of pachytene spermatocytes of about 80% purity based on the co-IF of SYCP3 and γH2AX (Fig. S6A-B).

**qRT-PCR of *miR-202-5p***

Total RNAs were isolated using Trizol regent (Invitrogen). Total RNAs were reverse-transcribed using miRcute miRNA First-strand cDNA Kit (TianGen) and the reactions were performed using miRcute miRNA qPCR Detection Kit (TianGen) in a Roche LightCycler 480 Real-Time PCR system (Primers: Table S3). The data were then analyzed using the comparative Ct method (ΔCt), with snoRNA234 (Snord70) RNA used as the internal control.

**Culture and meiosis initiation of SSCs**

Cells were maintained at 37°C under 5% $CO_2$ and tested negative for mycoplasma contamination. Establishment and maintenance of SSCs were performed following our previous reports (6). Briefly, testes from WT and KO mice at 6-7 days postpartum were digested with collagenase I and DNase I. The small seminiferous tubule fragments were plated on dishes in DMEM medium containing 10% FBS and SSCs were collected with several pipetting after 24 hours. Then, the SSCs were cultured on mitomycin C-inactivated mouse embryonic fibroblast feeder cells.

SSCs were digested with Accutase (Gibco) for 5 minutes, resuspended in DMEM medium containing 10% FBS and plated on a dish. After 30 minutes, MEF feeder cells but not SSCs attached to the dish bottom firmly. Floating SSCs were collected and plated to a plate coated with laminin (Sigma-Aldrich). Twenty-four hours later, cells were pre-treated with NC or Separase siRNA (CGUGUCCCUUUCACCAUAUUU) using Lipofectamine RNAiMAX reagent. After 10 hours, cells were treated by RA of 100 nM for 1 day, to induce meiosis initiation. Four days after induction, the cells entered meiosis prophase I and were harvested for characterization (Fig. 6F).

#### Supplementary Figures

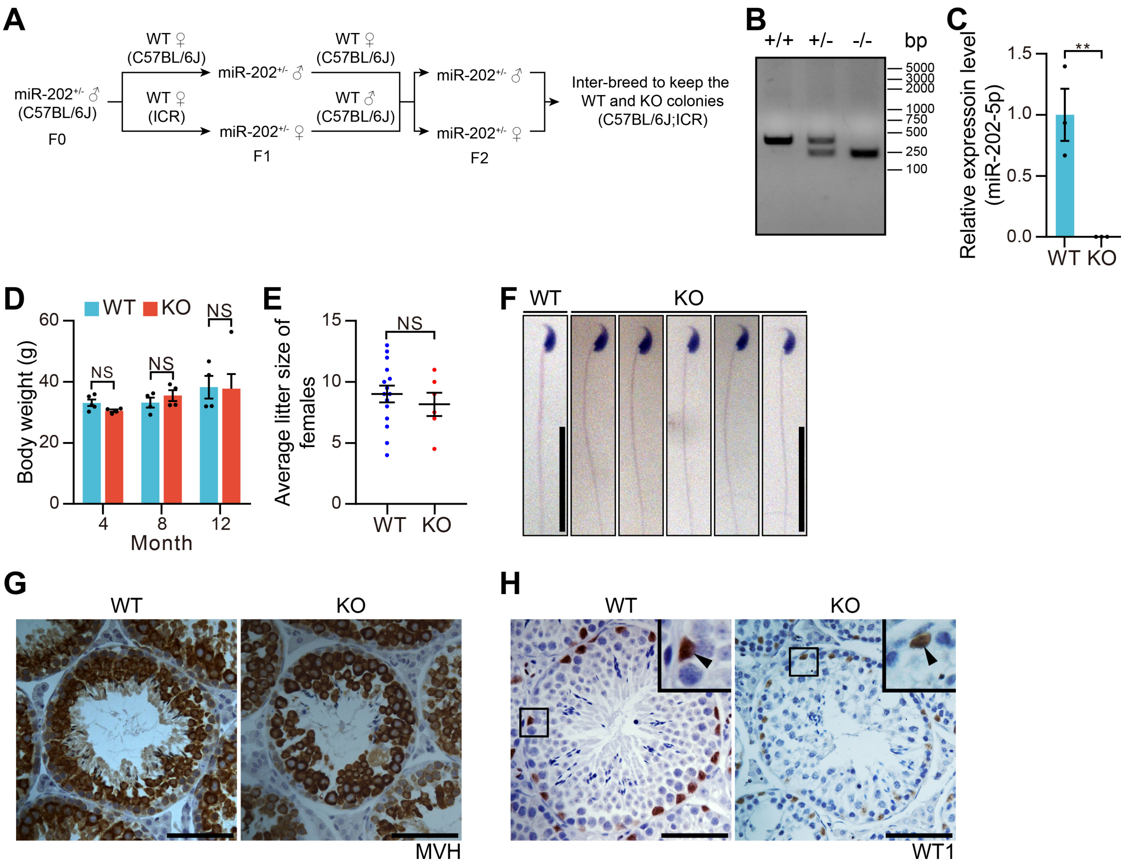


**Fig. S1. Construction and characterization of *miR-202* KO mice.** (**A**) Breeding strategy to construct WT and KO male mice under a C57BL/6J;ICR mixed background. (**B** and **C**) Deletion of *miR-202* in mice was verified by genomic PCR (B) and qRT-PCR (C). (**D**) Quantitative comparison of body weight between WT and KO mice. (**E**) Average litter size of WT or KO females. Each female mated with three WT males. Plugged WT or KO females were counted for litter size and each point represents the average litter size for one female. (**F**) Morphology analysis of sperm in adult WT and KO male mice by HE staining. (**G** and **H**) Immunohistochemistry of sections from adult WT and KO male mice for germ cell marker MVH (G) and Sertoli cell marker WT1 (H). Insets show high magnification images of boxed regions. Arrowheads indicate WT1$^{+}$

Sertoli cells. All values are shown as mean ± SEM. **$p < 0.01$. NS, not significant. Scale bars: 50 µm (G, H) and 20 µm (F).

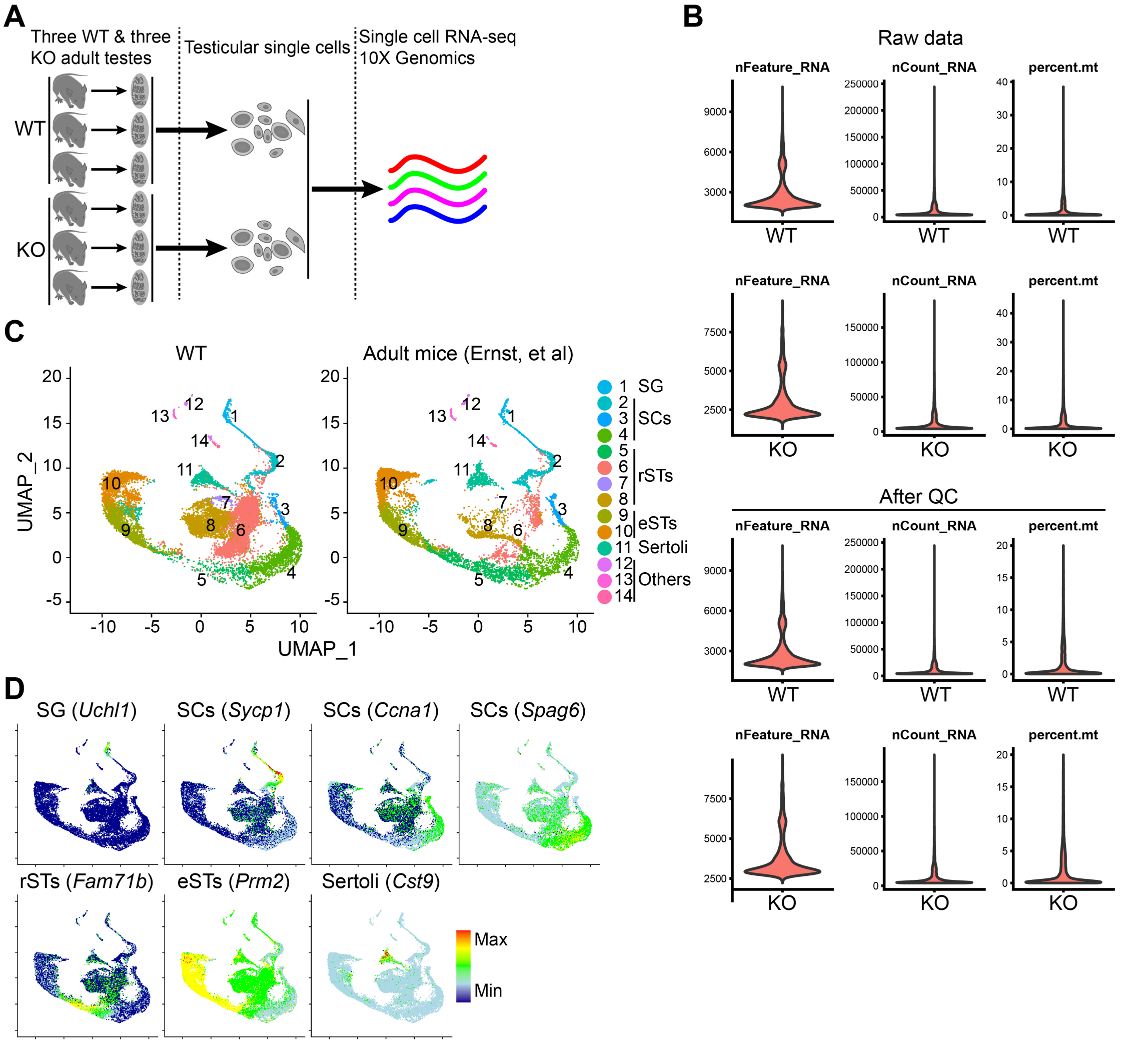


**Fig. S2. Single-cell RNA-seq of adult WT and KO testicular cells by 10x Genomics.** (**A**) Schematic illustration of the workflow for scRNA-seq analysis. Three WT testes and three KO testes were used for 10x Genomics single-cell RNA-seq. (**B**) Quality control (QC) metrics of raw and post-QC data for WT and KO. (**C**) UMAP and clustering analysis of single-cell transcriptome data from our WT mice and adult C57BL/6J mice published by Ernst, *et al*. Each dot represents a single-cell and cell clusters are distinguished by colors. SG, spermatogonia; SCs, spermatocytes; rSTs, round spermatids;

eSTs, elongating spermatids. (**D**) Gene expression patterns of selected cell marker genes projected on the combined UMAP plot in (C).

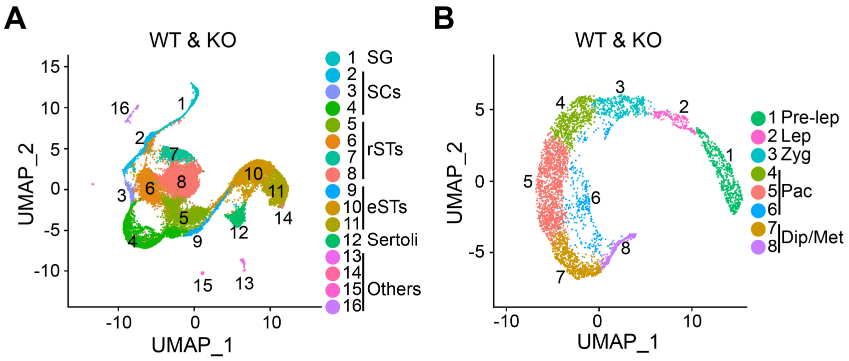


**Fig. S3. Total cell clustering and spermatocyte re-clustering for combined WT and KO.** (**A**) UMAP and clustering analysis of single-cell transcriptome data from combined testicular cells. Each dot represents a single-cell and cell clusters are distinguished by colors. SG, spermatogonia; SCs, spermatocytes; rSTs, round spermatids; eSTs, elongating spermatids. (**B**) UMAP and re-clustering analysis of single-cell transcriptome data from combined SCs of cluster 2, 3 and 4 in (A). Each dot represents a single-cell and cell clusters are distinguished by colors. Pre-lep, pre-leptonema; Lep, leptonema; Zyg, zygonema; Pac, pachynema; Dip/Met, Diplonema/metaphase.

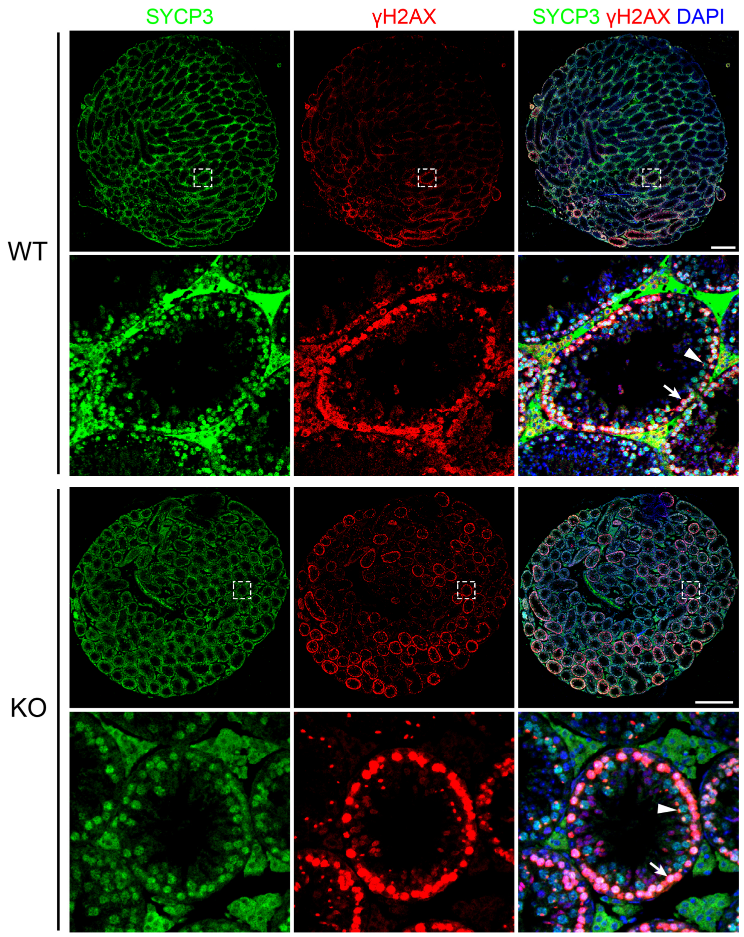


**Fig. S4. Images of whole testis sections from adult WT and KO mice.** Sections were immunostained for SYCP3 and γH2AX. The lower panels of WT and KO images are magnified tubules indicated by dashed areas, which contain both diffuse and dot-like γH2AX staining. Arrows and arrowheads indicate diffuse and dot-like γH2AX staining, respectively. Scale bars, 500 μm.

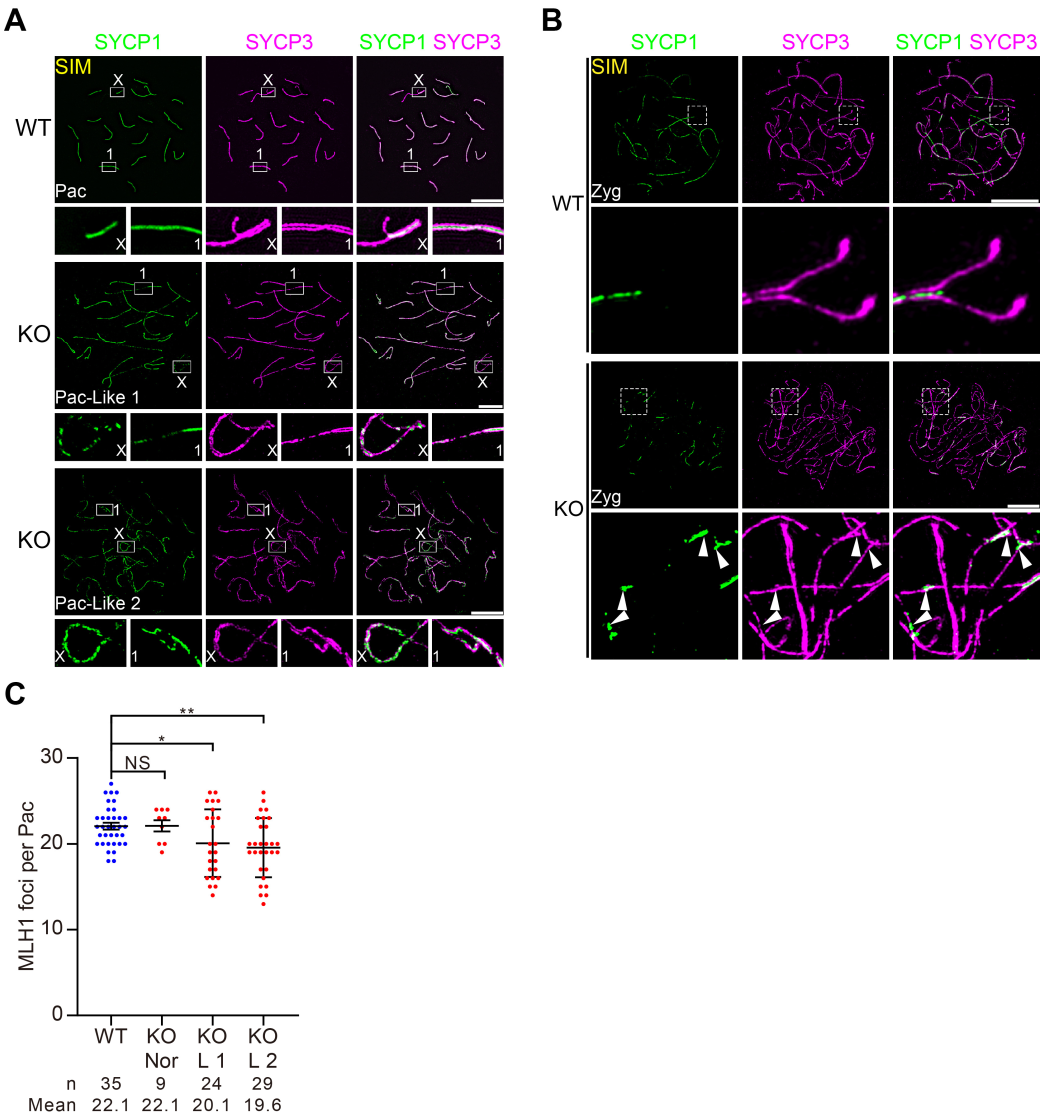


**Fig. S5. Loss of *miR-202* causes inter-sister synapsis and impaired crossover formation.** (**A**) Representative structured illumination microscopy (SIM) images of WT and KO Pac. Nuclear-spread SCs from WT and KO mice were immunostained for SYCP3 and SYCP1. Enlargements of boxes are shown below the respective full nucleus images, where "X" marks the XY body regions. (**B**) Representative SIM images of WT and KO Zyg. Nuclear-spread SCs from WT and KO mice were immunostained for SYCP3 and SYCP1. Enlargements of dashed regions are shown below the respective full nucleus images. Arrowheads represent the asynapsed chromosomes with SYCP1 signals, indicating inter-sister synapsis in KO. (**C**) Quantification of MLH1 focus numbers per Pac. At least three mice of each group were counted. Nor, normal Pac; L 1, Pac-like 1; L 2, Pac-like 2. All values are shown as mean ± SEM. $*p < 0.05$, $**p < 0.01$. NS, not significant. Scale bars: 10 μm.

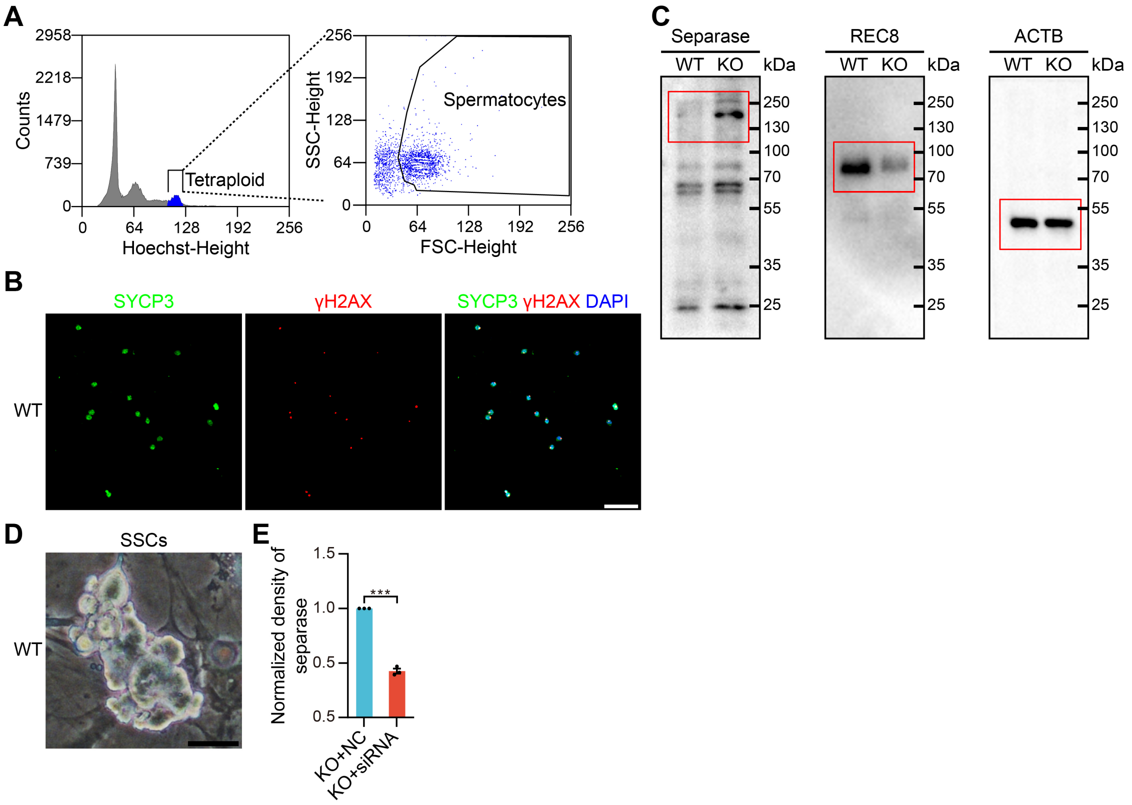


**Fig. S6. Isolation of SCs and culture of SSCs.** (**A**) Isolation of SCs from adult WT and KO testes after Hoechst staining by FACS. Tetraploid are gated by Hoechst staining, spermatocytes of which are further sorted. (**B**) Immunostaining evaluation of sorted spermatocytes for SYCP3 and γH2AX. (**C**) Original un-cropped Western blots shown in Fig. 5F (red boxed regions). (**D**) Typical images of cultured SSCs from WT mice. (**E**) Normalized density of separase in Fig. 6G indicates the efficient knockdown efficiency of seprase siRNA. All values are shown as mean ± SEM. ***$p < 0.001$. Scale bars: 100 µm (B) and 25 µm (D).

#### Supplementary Tables

**Table S1.** Quality control metrics of raw data and postQC data, and summary of cell clustering and re-clustering.

**Table S2.** Differentially expressed genes (DEGs) between WT and KO in spermatocytes, and biological process of GO terms enriched in DEGs.

**Table S3.** List of primers.

| Target | Forward | Reverse | Application |
|---|---|---|---|
| *miR-202* locus | AAGATCCGCTTGCGTAGGAA | CCAAGCTTAGTGGGGCTCTTT | Genomic PCR |
| *snoRNA234* | GTGATTTAACAAAAATTCGTCACTACCACTGAG | | qPCR |
| *miR-202-5p* | CCGGCGCGTTCCTATGCATATACTTCTTT | | qPCR |
| Separase | GCTGTCTTGTCTATGGTAGAAGC | TCATACAAAACACCAGGGAAAATC | Amplification of mRNA 3' UTR |

**Table S4.** List of antibodies.

| Antibody | Host | Company | Dilution | Cat # |
|---|---|---|---|---|
| SYCP3 | Mouse | Abcam | IF: 1:200 | ab97672 |
| SYCP3 | Rabbit | Abcam | IF: 1:200 | ab15093 |
| MVH | Rabbit | Abcam | IHC: 1:200 | ab13840 |
| WT1 | Rabbit | Epitomics | IHC: 1:200 | 2797-1 |
| γH2AX | Rabbit | CST | IF: 1:200 | 9718 |
| SYCP1 | Rabbit | Abcam | IF: 1:200 | ab175191 |
| REC8 | Rabbit | Abcam | IF: 1:200<br>WB: 1:2000 | ab192241 |
| RAD51 | Rabbit | Calbiochem | IF: 1:200 | PC130 |
| DMC1 | Rabbit | Santa Cruz | IF: 1:100 | sc-22768 |
| MLH1 | Mouse | BD Pharmingen | IF: 1:50 | 550838 |
| Separase | Rabbit | Abcam | IF: 1:200<br>WB: 1:1000 | ab3762 |
| ACTB | Mouse | Biodragon | WB: 1:5000 | B1029 |
| H1t | Guinea Pig | Kindly provided by M.A. Handel, Jackson Laboratory | IF: 1:500 | NA |
